# The bacterial iron sensor IdeR distinguishes DNA sequences by applying stress to non-cognate DNA

**DOI:** 10.64898/2026.10.05.756664

**Authors:** Suvankar Ghosh, Oksana Koshla, Dirk Maurer, Stijn Tieleman, Johan Åqvist, Julia J. Griese, Priyadarshi Satpati

## Abstract

The transcription factor IdeR regulates iron homeostasis in many gram-positive bacteria. It primarily functions as a repressor of iron uptake genes by binding to their promoters and blocking RNA polymerase access. IdeR recognizes a highly conserved DNA sequence consisting of four palindromic 5’-C_1_C_2_T_3_A_4_A_5_ repeats. For target site recognition, IdeR requires a thymine in position 3 of the repeat and at least one cytosine in the first two positions, while the identity of the bases in positions 4 and 5 is less important. We have previously shown that IdeR recognizes its target sequences through indirect or shape readout, rather than direct interactions with the bases. However, the mechanism of indirect recognition is unknown. Here we employed molecular dynamics simulations, X-ray crystallography, and DNA-binding assays to investigate the thermodynamic principles underlying accurate pyrimidine recognition by IdeR. We found that the recognition helix of IdeR forms a differential dielectric cleft that accommodates the C-H edge of the 5’-C_1_C_2_T_3_ sequence, with a polar pocket for the first two bases and a non-polar pocket for the third. The local dielectric environment of IdeR applies strain to non-cognate DNA, leading to non-productive binding and thereby ensuring the fidelity of target DNA reading.

**GRAPHICAL ABSTRACT:** 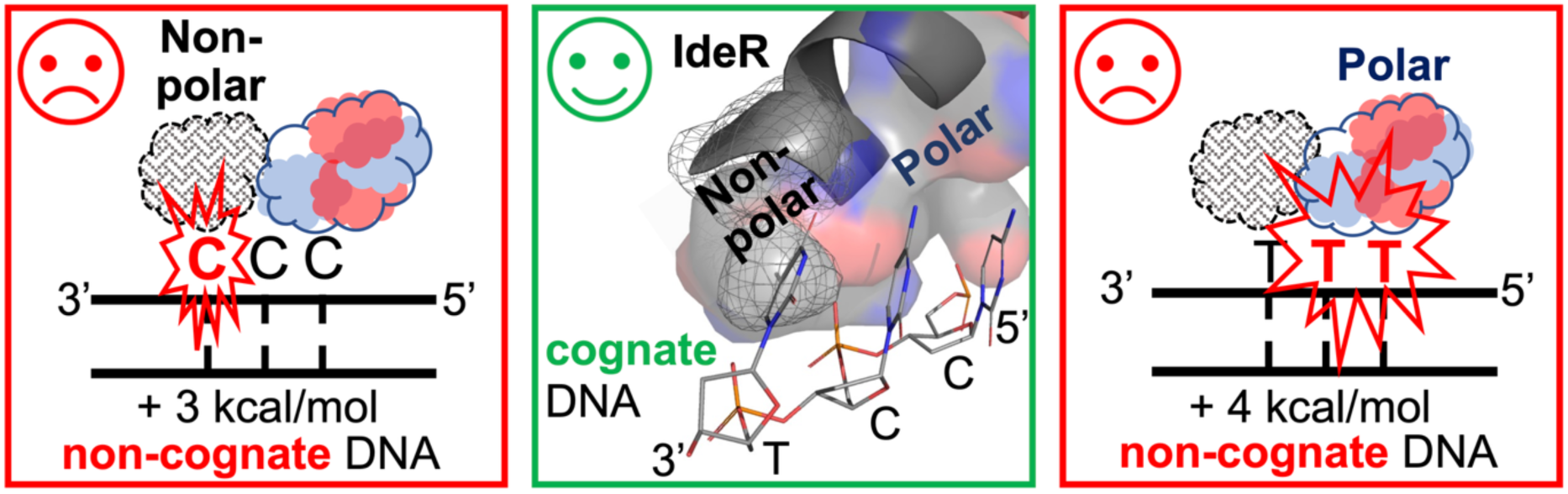

Alt text: Graphical abstract illustrating IdeR recognition of cognate DNA through a differential dielectric environment, with polar and non-polar pockets accommodating C_1_C_2_T_3_, while non-cognate sequences incur energetic penalties and binding strain.

## INTRODUCTION

DNA-binding proteins employ two different mechanisms to recognize their DNA targets. They can recognize specific base sequences by reading the chemical signatures of the DNA bases, in a mechanism termed direct or base readout, but they can also identify their binding sites by the sequence-dependent shape of the DNA backbone. This mechanism is called indirect or shape readout (1, 2). DNA-binding proteins can read both global and local DNA shape features which deviate from the ideal B-DNA conformation, including its sequence-dependent conformational flexibility/deformability and electrostatic potential (1–3). Some DNA-binding proteins utilize only one or the other, while many use a combination of base and shape recognition mechanisms (1, 2). Although it is well known that eukaryotic DNA-binding proteins commonly employ shape readout, the contribution of this mechanism to target recognition by prokaryotic transcription factors is still underexplored (1, 3–10).

Bacteria utilize a set of metal-responsive transcriptional regulators to adjust cellular metabolism to the metal availability in their environment. In gram-positive bacteria with high GC content, a transcription factor from the IdeR/DtxR (iron-dependent repressor/Diphtheria toxin repressor) family controls iron homeostasis (11, 12). This transcription factor contains an N-terminal winged helix-turn-helix domain, a helical dimerization domain, and a C-terminal SH3-like domain (12, 13). All three domains contribute to the two metal-binding sites of each subunit. When the intracellular iron level is sufficient, and the IdeR metal-binding sites are fully occupied by Fe^2+^, IdeR binds to its DNA target sites and prevents RNA polymerase from accessing the promoters of its target genes, thereby repressing transcription of primarily iron uptake genes (13).

We previously dissected the DNA recognition mechanism of IdeR from the actinomycete *Saccharopolyspora erythraea* (10). IdeR recognizes a highly conserved 19-bp palindromic sequence consisting of four 5’-C_1_C_2_T_3_A_4_A_5_ repeats (TTAGGTTAGSCTAACCTAA; where S = G or C, i.e. G in the forward strand and C in the reverse strand) (10, 14–16). Two IdeR dimers bind cooperatively to its canonical DNA target sites, with the dimers sitting on opposite sides of the DNA double helix, one dimer contacting CCTAA repeats 1 and 3 and the other repeats 2 and 4 (10, 14, 17–22) (Fig. 1A).

**Figure 1.**
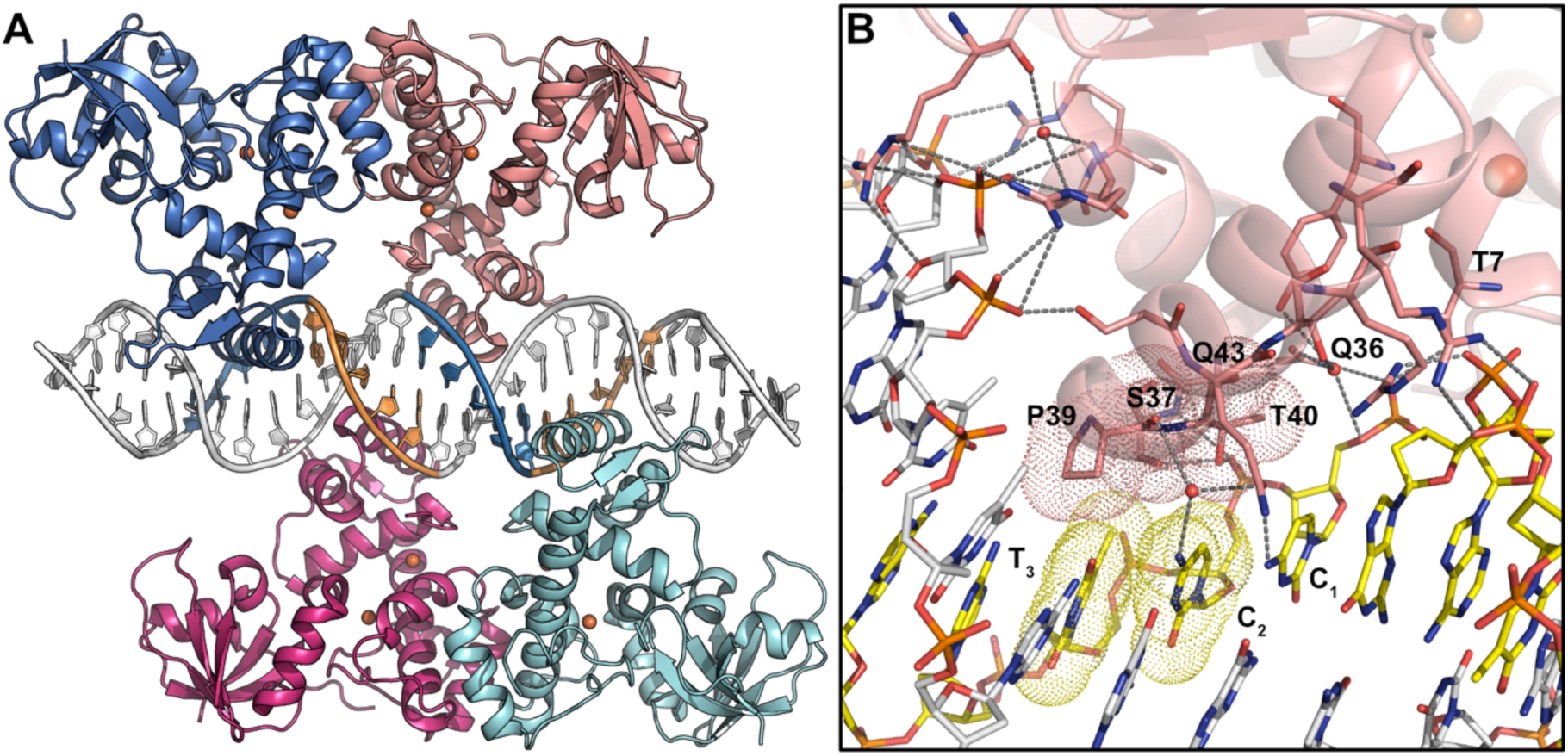
Crystal structure of Fe^2+^-activated IdeR in complex with the consensus DNA-binding sequence (PDB 7B20) (10). (**A**) Overall structure of the complex in cartoon representation, with IdeR colored by subunits and the DNA repeats interacting with IdeR colored blue (repeats 1 and 3) and orange (repeats 2 and 4). The Fe^2+^ ions are shown as orange spheres. (**B**) Detailed view of the IdeR–DNA interactions, illustrated on IdeR subunit B. Sticks are shown colored by element with salmon carbon atoms for the protein, while the two DNA strands are shown with yellow and white carbon atoms, respectively. Hydrogen bonds and salt bridges are indicated by grey dashed lines. Van der Waals interactions are indicated by the dotted van der Waals surfaces of IdeR residues Ser37, Pro39 and Thr40 as well as the DNA bases C_2_ and T_3_. Gln43 forms direct and water-mediated hydrogen bonds with DNA bases C_1_ and C_2_, respectively. Other polar and charged residues in the recognition helix, the N-terminal helix and the wing of the winged helix-turn-helix motif form hydrogen bonds and salt bridges with the DNA backbone on either side of the major groove into which the recognition helix inserts. Residues mentioned in the text are labelled. Alt text: Crystal structure of Fe^2+^-activated IdeR bound to consensus DNA, showing the overall protein–DNA complex (A) and close-up of the protein–DNA interactions with hydrogen-bond, salt-bridge, and van der Waals interactions (B).

By systematically mutating the bases of the CCTAA motif and the amino acid residues in the IdeR recognition helix which contact these bases, we demonstrated that IdeR uses a shape readout mechanism to identify its target sites. We showed that IdeR requires a thymine in position 3 of the repeat and a cytosine in at least one of the first two positions, while the identity of bases 4 and 5 is less important, although not irrelevant for target recognition (10). IdeR makes very few contacts with the bases in the recognition motif, and only one direct hydrogen bond is observed: Gln43 in the recognition helix forms a hydrogen bond with the cytosine (or guanine) base in position 1, as well as a water-mediated hydrogen bond with the cytosine in position 2 of the repeat (Fig. 1B). However, both of these interactions were shown to be expendable for recognition. Additionally, Ser37 and Pro39 form van der Waals interactions with the thymine in position 3, and Thr40 also interacts with the cytosine backbone in position 2, while no direct interactions are observed with the bases in positions 4 and 5. We demonstrated that Pro39 is important, although not essential, for sequence recognition.

The current view rules out a base readout mechanism for IdeR target recognition and strongly supports a shape readout mechanism (10). However, our earlier experiments did not address how exactly shape recognition is achieved. Therefore, we turned to Molecular Dynamics (MD) simulations to further investigate this intriguing and fundamental question. Together with crystallographic data and DNA-binding assays, we show that the dielectric environment of the IdeR recognition helix favors interactions only with target sequences, but exerts stress on non-target sequences, thereby ensuring accurate DNA recognition.

## MATERIAL AND METHODS

### Molecular Dynamics Setup and Simulation

The crystal structure of the Fe^2+^-activated IdeR–consensus-DNA complex (PDB 7B20) (10) was used. One chain of the biomolecule (chain B) was positioned at the centre of an equilibrated water box with dimensions 10 × 10 × 10 nm³. Counterions (36 Na^+^) were added to neutralize the system. Biomolecular interactions were described using the CHARMM36m force field (23). The water molecules were modelled using the TIP3P model (24). The solvated simulation box underwent energy minimization for 50,000 steps using the steepest descent algorithm (25), followed by equilibration in two phases: first NVT (200 ps) and then NPT (200 ps) ensembles at 310 K and 1 bar. Production simulations (NPT at 310 K and 1 bar) were carried out for 500 ns using a 2 fs time step, with long-range electrostatics treated using the particle-mesh Ewald method (26) and covalent bonds involving hydrogens constrained using LINCS (27). Multiple independent molecular dynamics simulations were performed using GROMACS (v2019.6) (28–30). The production dynamics were used for structural analysis, and the final structures from the production runs were subjected to alchemical free energy simulations.

### Molecular Dynamics Free Energy Calculations

Alchemical free energy calculations were conducted to assess how a DNA base-pair mutation affects the IdeR binding affinity. The mutation was modelled using a dual-topology approach, which involved alchemically transforming the cognate DNA base pair into a non-cognate base pair during the simulation. This transformation was performed across a series of 51 λ-windows, with λ values ranging from 0 to 1. Each window was equilibrated for a specified time (200 ps), followed by a production simulation lasting 5 ns. To avoid singularities during the transformation, soft-core potentials were utilized. The alchemical transformation was carried out both in the presence and absence of the IdeR protein. An appropriate thermodynamic cycle (Fig. 2) was employed to calculate the difference in IdeR binding affinity between the cognate and non-cognate DNA. A similar approach was used to estimate the change in DNA-binding affinity resulting from IdeR mutations. Free energy differences were calculated using the Bennett acceptance ratio (BAR) method (31), as implemented in GROMACS (Supplementary Table S1).

**Figure 2.**
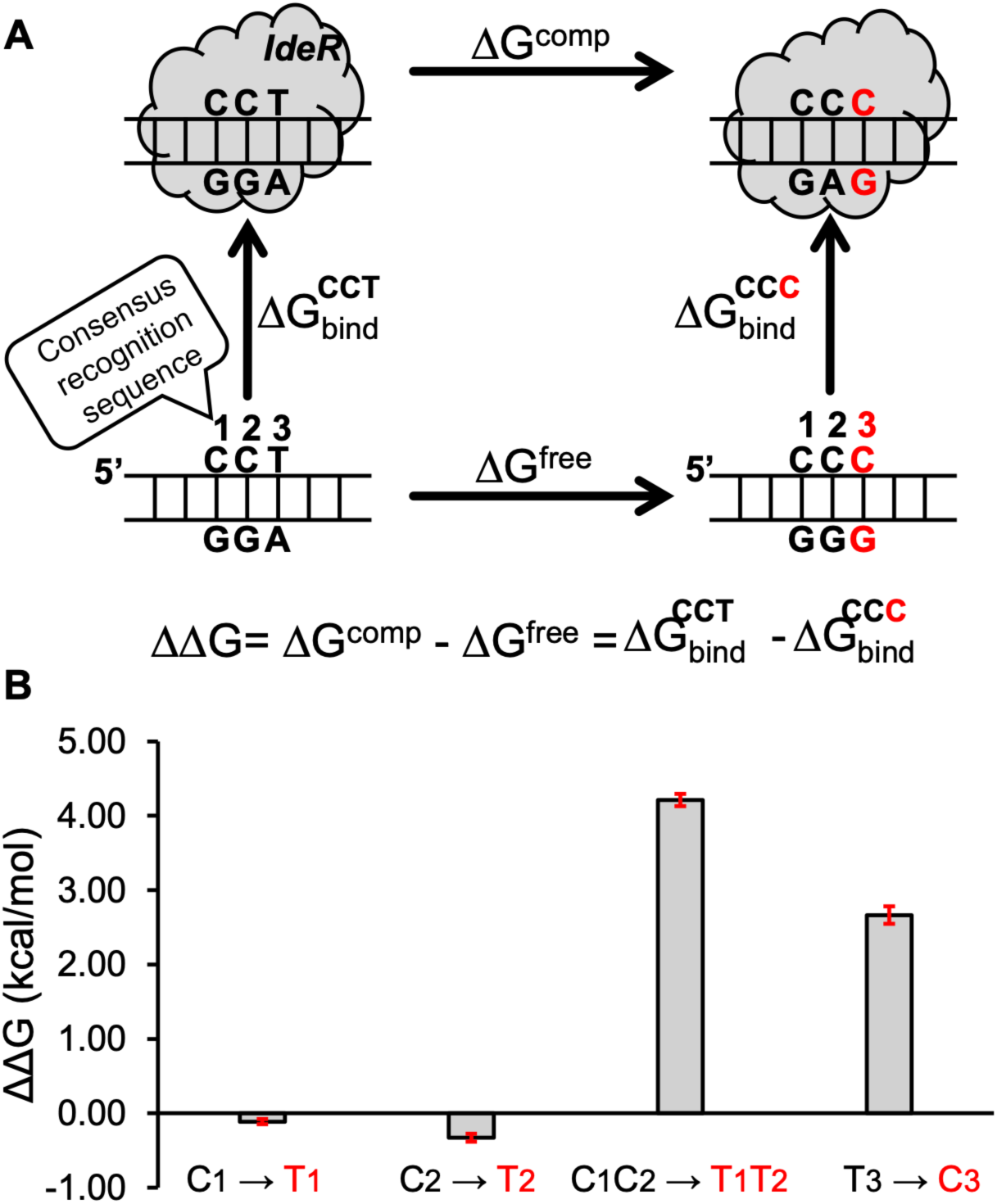
(**A**) Thermodynamic cycle and IdeR–DNA binding. Vertical arrows show DNA binding to IdeR, while horizontal arrows depict the transformation of cognate DNA base pairs (T-A) into non-cognate form (C-G), either bound to IdeR (upper arm) or free in water. The relative binding free energy is calculated as ΔΔG = ΔG^comp^ – ΔG^free^. (**B**) The ΔΔG plot shows the IdeR binding affinity for non-cognate DNA sequences compared to consensus DNA, with error bars indicating the standard error of the mean (SEM). Alt text: Thermodynamic cycle illustrating IdeR binding to consensus and non-cognate DNA sequences (A), with calculated relative binding free energies for four base-pair substitutions shown as a bar graph (B).

MM/PBSA calculations were conducted using a python script (32) to estimate the absolute DNA binding affinity (ΔG_bind_) of the wild-type and P39G-variant IdeR. The binding free energy of the DNA–protein complex was calculated from snapshots extracted from the production trajectory. A total of 1000 frames were extracted at regular intervals from the last 100 ns of the production run and used for the calculation. The binding free energy was computed as the difference between the free energy of the complex and the sum of the free energies of the isolated protein and DNA. Molecular mechanics energy terms included van der Waals and electrostatic interactions, while solvation free energies were decomposed into polar contributions calculated using the Poisson–Boltzmann equation and nonpolar contributions estimated from the solvent-accessible surface area. The entropy was estimated from the interaction entropy (IE) method (33). MM/PBSA calculations were performed using two solute dielectric constants of ε = 1 and 5, while keeping all other default parameters (Supplementary Table S1).

### Protein production and purification

*S. erythraea* IdeR^WT^ was recombinantly produced in *E. coli* BL21(DE3) (Novagen) with an N-terminal, TEV-cleavable hexahistidine tag and purified via immobilized metal ion affinity chromatography (IMAC), removal of the hexahistidine tag by incubation with TEV protease, and reverse IMAC, as previously described (10). The purified protein was concentrated to a protein concentration of ∼1.75 mM (45 mg/ml) in 25 mM MES pH 6.0, 450 mM NaCl, 10% (v/v) glycerol. Protein concentration was determined using a calculated extinction coefficient at 280 nm of 15.47 mM^-1^ cm^-1^ for the IdeR monomer (34). The protein was aliquoted, flash-frozen in liquid nitrogen and stored at-70°C until further use.

### Preparation of double-stranded DNA

Forward and reverse DNA oligonucleotides containing the target sequences designed for DNA-binding analysis and crystallization studies were obtained from Eurofins with or without a 5’ FAM fluorescent label (Supplementary Table S2). To prepare double-stranded DNA, each forward and reverse oligonucleotide sample pair was resuspended in DNA buffer (40 mM Tris-HCl pH 7.4, 100 mM NaCl, 10 mM MgSO_4_) and mixed to a final concentration of 1 mM (for co-crystallization with IdeR) or 10 µM (for EMSA) each in DNA buffer. The mixtures were then heated at 95°C for 5 min and slowly cooled down to 4°C in a thermal cycler.

### Electrophoretic mobility shift assays (EMSAs)

EMSAs were performed after mixing the fluorescently-labelled double-stranded DNA samples at 30 nM with 30 µM (for the standard IdeR concentration range) or 0.1 mM CoCl_2_ (for the 10x higher IdeR concentration range) and IdeR^WT^ at different concentrations (standard range 15 nM – 2.25 µM dimer; 10x higher range 0.15 – 22.5 µM dimer). All mixtures were prepared in TAKA buffer (15 mM Tris-acetate pH 7.3, 4 mM potassium-acetate), containing glycerol at a final concentration of 10% and Poly(dI-dC) (Poly(deoxyinosinic-deoxycytidylic) acid; Thermo Fisher Scientific) as competitor DNA at a final concentration of 17 ng/µl. The samples were incubated at room temperature for 20 min before loading on a 4% native polyacrylamide gel. Each EMSA was run in TAKA buffer at 4°C for 55 min at 10 mA/gel. Fluorescence of the unbound DNA and the DNA bound to IdeR was then observed in a BioRad ChemiDoc MP Imager using the proper filter for the FAM label (530/28 Filter). EMSAs were performed at least twice and with different protein batches to ensure reproducibility. As positive control, EMSAs were performed with the full consensus IdeR recognition sequence to ensure that the protein batch used was active.

### Crystallization and data collection

Crystallization conditions were screened using commercial kits (Molecular Dimensions) in sitting-drop vapor diffusion setups at 20°C using a Mosquito^®^ Crystal liquid handling robot (SPT Labtech), followed by optimization of the identified condition. To obtain the structure of IdeR^WT^ complexed with cobalt and the Consensus_C_2_-T_2_ DNA, IdeR^WT^ at a final concentration of 0.47 mM (12 mg/ml) was mixed with 1 mM CoCl_2_ and 150 µM double-stranded DNA (Supplementary Table S2) in buffer containing 25 mM MES pH 6.0, 50 mM NaCl and 10% (v/v) glycerol, to reduce the salt concentration to ∼160 mM in the final sample. The sample was incubated at room temperature for 30 min. This solution was then mixed with crystallization solution containing 20-32.5% (w/v) PEG 3350, 200 mM sodium acetate and 100 mM Bis-Tris-HCl at pH 5.0, 5.5 or 6.0 in a sitting-drop vapor diffusion experiment at 20°C using the Mosquito^®^ Crystal robot. The total drop volume was 600 nl and the protein volume was 150 nl or 300 nl. Crystals were harvested after addition of 20% glycerol to the mother liquor as cryoprotectant and flash-cooled in liquid nitrogen. A dataset of the IdeR^WT^ complex with Co^2+^ and Consensus_C_2_-T_2_ DNA was obtained from a crystal grown in 22.5% (w/v) PEG 3350, 200 mM sodium acetate and 100 mM Bis-Tris-HCl at pH 5.0. The dataset was collected at 100 K at beamline I24 of Diamond Light Source (Didcot, United Kingdom) (Table 1).

**Table 1.**
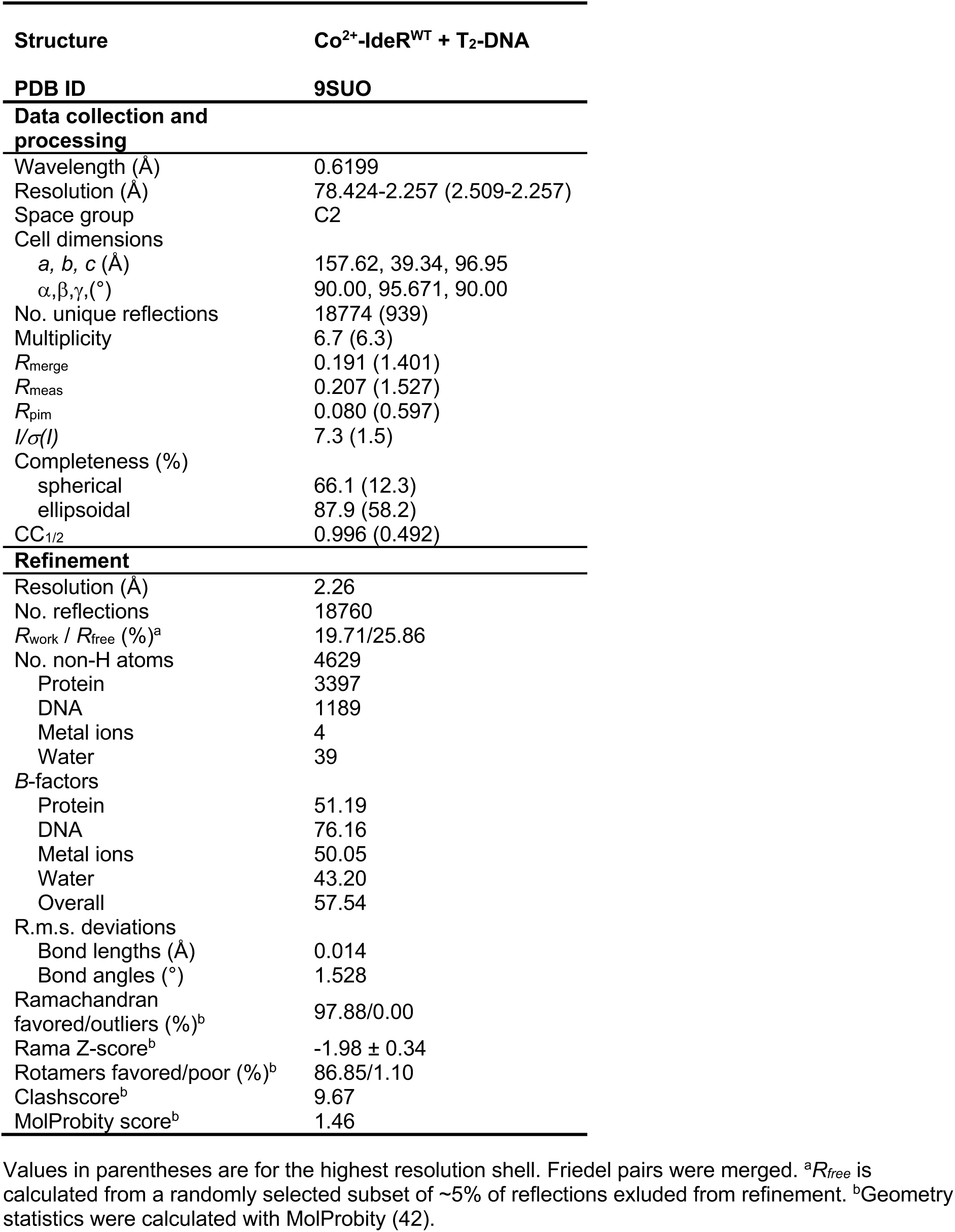
Crystallographic data and refinement statistics.

### Structure determination, model building and refinement

Two 180° sweeps of the same crystal were processed with XDS and merged with XSCALE (35). Since diffraction was severely anisotropic, an elliptical diffraction cut-off was chosen using the STARANISO server (http://staraniso.globalphasing.org/cgi-bin/staraniso.cgi) based on the criterion that the local I/σ(I) ≥ 1.20. The IdeR^WT^-Consensus_C_2_-T_2_ complex crystallized in space group C2 with one IdeR dimer and one DNA strand in the asymmetric unit, so that the complete complex containing a DNA duplex and two IdeR dimers is created by crystallographic symmetry. This is possible because the DNA sequence is almost perfectly palindromic, with forward and reverse strands differing only in the central base of the palindrome (G or C) and the 5 bases at either end of the recognition sequence which do not interact with IdeR (3 of which are visible in the crystal structure; Supplementary Table S2), so that the protein does not discriminate between the two possible orientations of the DNA double strand (10). The structure was solved by molecular replacement using PHASER (36) in CCP4 (37) and a search model containing one IdeR dimer and one DNA strand, derived from the Fe^2+^-activated complex of IdeR with consensus DNA (PDB 7B20) (10). Refinement was carried out with REFMAC5 (38) and Phenix.refine (39, 40) and iterated with rebuilding in Coot (41). Two superimposed DNA strands at 50% occupancy each were built and treated as a constrained group during occupancy refinement. Refinement included bulk solvent corrections, individual atomic coordinate and isotropic *B* factor refinement as well as occupancy refinement for alternate conformations. Riding hydrogens were used during refinement. Solvent molecules were added with REFMAC5 and Phenix.refine and manually curated. Final refinement was carried out in Phenix.refine and included metal–ligand bond length restraints, TLS and torsion angle NCS restraints. The structure was validated using MolProbity (http://molprobity.biochem.duke.edu/) (42). Data and refinement statistics are given in Table 1. All figures were prepared with PyMOL (version 3.1.8; Schrödinger, LLC). The DNA conformation in the crystal structure was analyzed with DSSR (43), and protein–DNA interactions were analyzed with DNAproDB (44).

## RESULTS

### Structure and dynamics: insights from molecular dynamics simulations

MD simulations effectively reproduced the crystal structure of the IdeR–C_1_C_2_T_3_A_4_A_5_ complex (Fig. 1A), as evidenced by a low root-mean-square deviation (RMSD) (Supplementary Fig. S1A). The plateau observed in the RMSD plot indicates that the simulations achieved structural convergence and the relevant conformations were adequately sampled during the simulations. During the trajectories, the recognition helix of IdeR occupied the major groove of the consensus recognition sequence of the target DNA. The non-specific interaction between the backbone phosphate of C_1_ and protein side chains Thr7 and Gln36 was preserved throughout the simulations (Fig. 1B, Supplementary Fig. S1). As observed in the crystal structure (10), the first six residues of the N-terminal helix of IdeR were unstructured (Supplementary Fig. S1), which is essential for DNA binding (45). The side chain of the Gln43 residue exhibited flexibility and accessed bulk water during the MD trajectories (see Q43movie.mpg at https://doi.org/10.5281/zenodo.22951796). This finding suggests that both the direct and water-mediated interactions between the side chain of Gln43 and the amine groups of C_1_ and C_2_ observed in the crystal structure (Fig. 1B) are unstable during the trajectories.

The final structures from the production dynamics were then subjected to alchemical free energy simulations to estimate the thermodynamics of DNA binding to IdeR.

### Thermodynamics of DNA recognition by wild-type IdeR

Using the final production MD structure of the IdeR–C_1_C_2_T_3_ complex as a template, we performed molecular dynamics free energy calculations to study the energetics of DNA recognition by the wild-type IdeR protein. These calculations determine the change in IdeR binding affinity (ΔΔG) caused by base-pair mutations in the triplet consensus recognition sequence (C_1_C_2_T_3_) of the DNA (Fig. 2A). The calculated ΔΔG provides a quantitative estimate of how strongly IdeR prefers the consensus (C_1_C_2_T_3_) DNA compared to non-consensus sequences.

The energetics (ΔΔG) associated with base-pair mutations in the consensus DNA sequence, specifically the first position (C_1_→T_1_), second position (C_2_→T_2_), and third position (T_3_→C_3_), are summarized in Fig. 2B and Supplementary Table S1. IdeR demonstrated strong specificity for the third position of the DNA. The non-cognate base (C_3_) in the third position is significantly disfavored by IdeR, by ΔΔG ∼3 kcal/mol, compared to the cognate T_3_-DNA complex. This results in a non-cognate complex population as low as 1 in 1000. In contrast, IdeR does not distinguish between cytosine and thymine at either the first or second position of the DNA sequence for binding (ΔΔG < 0.5 kcal/mol). However, a double mutation (C_1_C_2_→T_1_T_2_) in the DNA significantly disfavors IdeR binding, leading to a penalty of ∼4 kcal/mol.

In our earlier experiments (10), we used a 19-base-pair consensus DNA sequence containing 4 palindromic C_1_C_2_T_3_A_4_A_5_ repeats to create several non-cognate mutant DNA sequences (with mutations present in all 4 repeats). The affinity of IdeR for these mutant and consensus DNA sequences was assessed using electrophoretic mobility shift assays (EMSAs). The results showed that replacing cytosine with thymine at either the first or second position of the repeat did not significantly affect IdeR affinity. The apparent dissociation constants (*K_D_*) for IdeR binding to the first and second C→T DNA mutants were estimated to be ∼70 nM and ∼105 nM, respectively, as compared to ∼45 nM for the full consensus DNA. The marginal change in affinity relative to the full consensus DNA is accurately reflected by the small computed ΔΔG values of-0.5 kcal/mol (Fig. 2B).

However, when both cytosines (C_1_C_2_) were simultaneously mutated to thymine (T_1_T_2_) in the DNA, IdeR did not bind (10). Thus, a cytosine base in at least one of the first two positions of the repeat is necessary for IdeR recognition, which is validated by the large calculated ΔΔG of 4 kcal/mol for the double thymine C_1_C_2_→T_1_T_2_-mutant.

Our calculations estimated that DNA binding to IdeR is weakened by approximately 3 kcal/mol due to the T_3_→C_3_ mutation in the consensus DNA, which is corroborated by the experimental observation of no binding between IdeR and C_1_C_2_C_3_A_4_A_5_ non-cognate DNA (Fig. 3 and Supplementary Fig. S2). We also calculated the effect of the T_3_→G_3_ mutation on IdeR binding affinity and found that the mutation reduced binding affinity by 2 kcal/mol, consistent with our previous experiments showing no binding to the C_1_C_2_G_3_A_4_A_5_ repeat (10).

**Figure 3.**
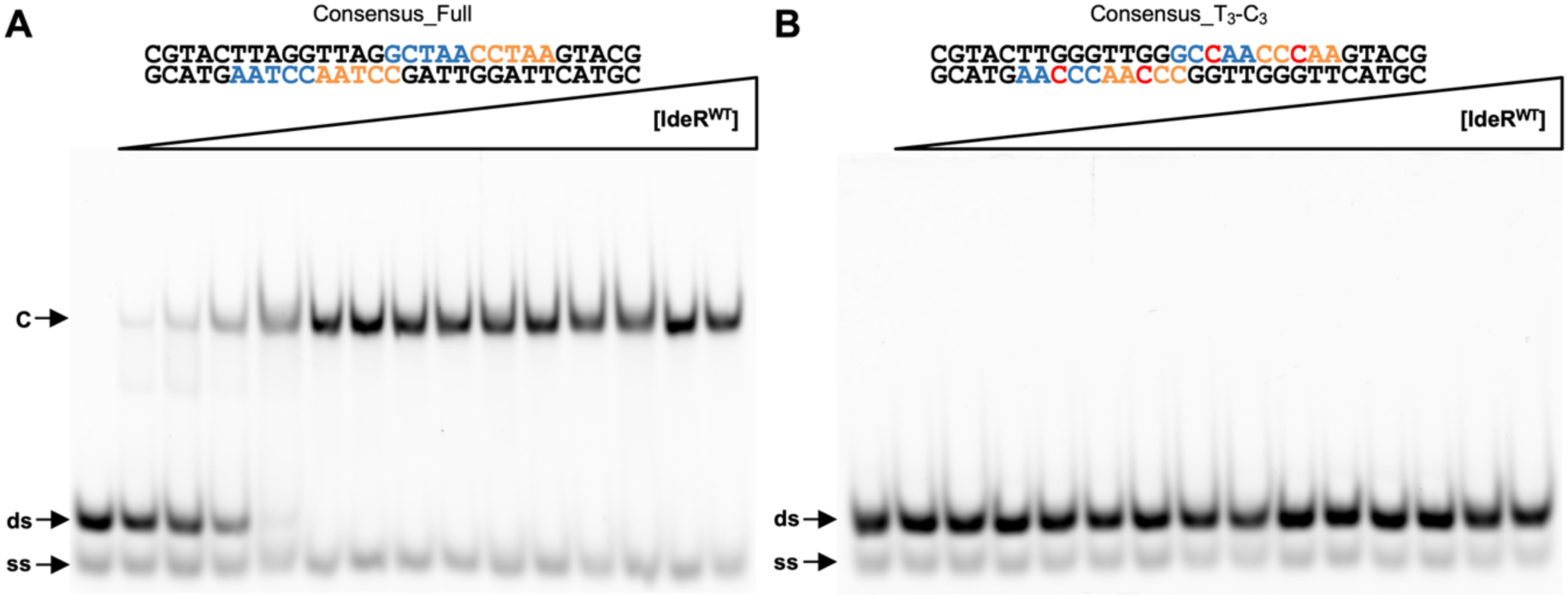
EMSA analysis of IdeR^WT^ binding to (**A**) the full consensus DNA recognition sequence, showing that the protein preparation used is active, and (**B**) the Consensus_T_3_-C_3_ DNA sequence, which demonstrates that IdeR does not recognize this sequence. IdeR was added in increasing concentrations (15 nM – 2.25 µM dimer) to 30 nM fluorescence-labeled double-stranded DNA probe in the presence of 30 µM Co^2+^ and competitor DNA. Protein– DNA complexes were resolved on a 4% Tris-acetate polyacrylamide gel. The left-most lane is a control reaction without protein. C, protein–dsDNA complex; ds, unbound double-stranded DNA probe; ss, non-hybridized single-stranded DNA. Alt text: EMSA showing increasing IdeR concentrations bind the full consensus DNA sequence (A), producing protein–DNA complexes, but do not bind the T_3_-C_3_ mutant sequence (B).

Thus, the calculated ΔΔG effectively reproduced our experimental observations of the effect of mutations in the DNA recognition sequence on IdeR binding affinity.

### The hydrophobic cushion of Pro39 of IdeR ensures selectivity for thymine at the third position of the DNA repeat

Although our previous data suggested that Pro39 is important for target recognition, its precise role could not be ascertained based on mutational data, crystal structures and DNA-binding assays (10).

The major groove of the recognition sequence (C_1_C_2_T_3_), particularly at the third position (T_3_), is desolvated by residue Pro39 in the IdeR recognition helix. The proline creates a hydrophobic cushion for the methyl group of the thymine at position 3 (Fig. 1B). Mutations at the third position (T_3_→C_3_: Fig. 2B; or T_3_→G_3_: Supplementary Table S1) increase the polarity of the major groove. As a result, the mutated third base pair (CG or GC) within the non-polar pocket deforms the DNA backbone in the IdeR complex. This deformation is observed in the MD structure (Fig. 4). The strain on the DNA backbone negatively impacts IdeR binding by increasing the free energy of the non-cognate IdeR–DNA complex. Therefore, it can be argued that the selectivity for thymine at the third position is maintained by the hydrophobic cushion provided by Pro39, as previously anticipated (21).

**Figure 4.**
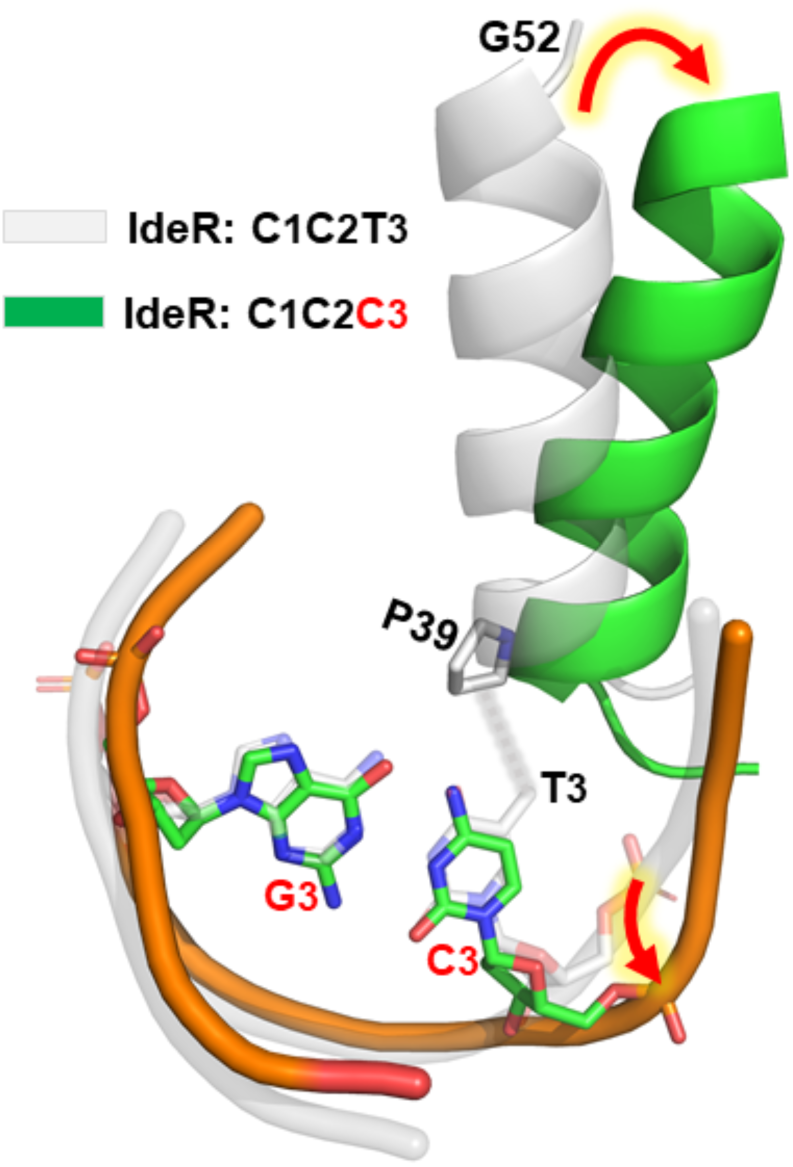
The mutation from T_3_ to C_3_ resulted in the loss of interaction represented by a broken, grey, glowing line between the 5-methyl group of T_3_ and the Pro39 residue of IdeR. This change altered the orientation of the recognition helix and introduced strain into the DNA, as indicated by the red arrows with a yellow glow. Alt text: Structural comparison of IdeR bound to C_1_C_2_T_3_ full consensus and C_1_C_2_C_3_ mutant DNA, showing loss of the T_3_ 5-methyl–Pro39 interaction, altered IdeR recognition-helix orientation, and DNA strain.

In EMSAs, the IdeR^P39G^ mutant showed a six-fold decrease in affinity for cognate DNA compared to IdeR^WT^, highlighting the importance of Pro39 in DNA binding (10). We conducted simulations of the IdeR^P39G^:C_1_C_2_T_3_ complex by replacing Pro39 with Gly39 in the template crystal structure. Weak protein–DNA interaction was evident in the MD trajectories due to the loss of the Pro39–T_3_-CH_3_ interaction. Estimation of absolute binding free energy using the faster and computationally less expensive MM/PBSA approach indicated that the P39G mutation in IdeR compromises cognate C_1_C_2_T_3_ binding by 3 kcal/mol (Supplementary Table S1). Thus, it is evident that Pro39 plays a key role in the recognition of the target DNA sequence.

### The Gln43 residue minimally affects IdeR binding to DNA

The crystal structure of the IdeR–C_1_C_2_T_3_A_4_A_5_ complex revealed only one base-specific hydrogen bond between the protein and the DNA, formed by Gln43 with C_1_, as well as a water-mediated hydrogen bond between the same residue and the C_2_ base (10), suggesting that Gln43 might play a role in DNA recognition. However, EMSAs showed that the DNA-binding affinity of IdeR^Q43A^ is not significantly different from that of IdeR^WT^ (10). We evaluated the impact of the Q43A mutation in IdeR on its consensus C_1_C_2_T_3_ binding affinity. Alchemical free energy calculations confirmed that the DNA binding affinity is unaffected by the Q43A mutation in IdeR (ΔΔG =-0.13 ± 0.15 kcal/mol, Supplementary Table S1). We also found that residue Gln43 was flexible and primarily interacted with bulk water in the MD trajectories (Supplementary Fig. S1). The structure of the IdeR–C_1_C_2_T_3_ complex was largely unaffected by the Q43A mutation, except that water molecules filled the space created by the mutation, in agreement with the crystal structure of this complex (10). Thus, Gln43 alone does not contribute significantly to DNA binding affinity.

### A polar pocket in the IdeR recognition helix selects for cytosine bases in the first two positions of the DNA repeat

Simulation showed that the interaction network of the IdeR–DNA complex was not affected by a single mutation at the first or second base pair of the DNA repeat (C_1_→T_1_ or C_2_→T_2_). The structures containing the respective mutation closely resemble the template crystal structure IdeR–C_1_C_2_T_3_ complex. To corroborate the simulation, we determined the structure of the IdeR–C_1_T_2_T_3_ complex by X-ray crystallography.

IdeR^WT^ was crystallized in complex with Co^2+^ and the DNA sequence containing the C_2_ →T_2_ mutation in every repeat. The structure of the complex was determined at a resolution of 2.26 Å in space group C2, containing 1 IdeR dimer and 1 DNA strand in the asymmetric unit. The DNA represents a superposition of the 29-bp forward and reverse strands, with approximately equal occupancy. This is possible because the protein does not distinguish between the two orientations of the DNA duplex, as its recognition sequence is palindromic and it does not interact with the non-palindromic flanking base pairs outside the recognition sequence. The complete assembly, consisting of 2 IdeR dimers and 1 DNA duplex, is obtained by applying crystallographic symmetry.

The crystal structure is essentially identical to our predicted structure. At first glance, the complex structure also appears nearly identical to that of IdeR complexes with the full consensus recognition sequence (10). However, when superimposing the IdeR–T_2_-DNA complex with the IdeR–consensus-DNA complex (PDB 7B20), it becomes apparent that one of the IdeR dimers in the T_2_-DNA complex is shifted with respect to the corresponding dimer in the consensus-DNA complex (Fig. 5A and Supplementary Fig. S3). This shift appears to result from differences in the structures of the respective DNA duplexes. The DNA backbone is significantly deformed around position 2 of the repeat in the T_2_-DNA complex compared to the C_2_-DNA complex (Fig. 5A). One IdeR dimer must therefore adopt a different orientation in relation to the dimer bound to the opposite face of the T_2_-DNA duplex to be able to access the major groove of this non-cognate DNA sequence.

**Figure 5.**
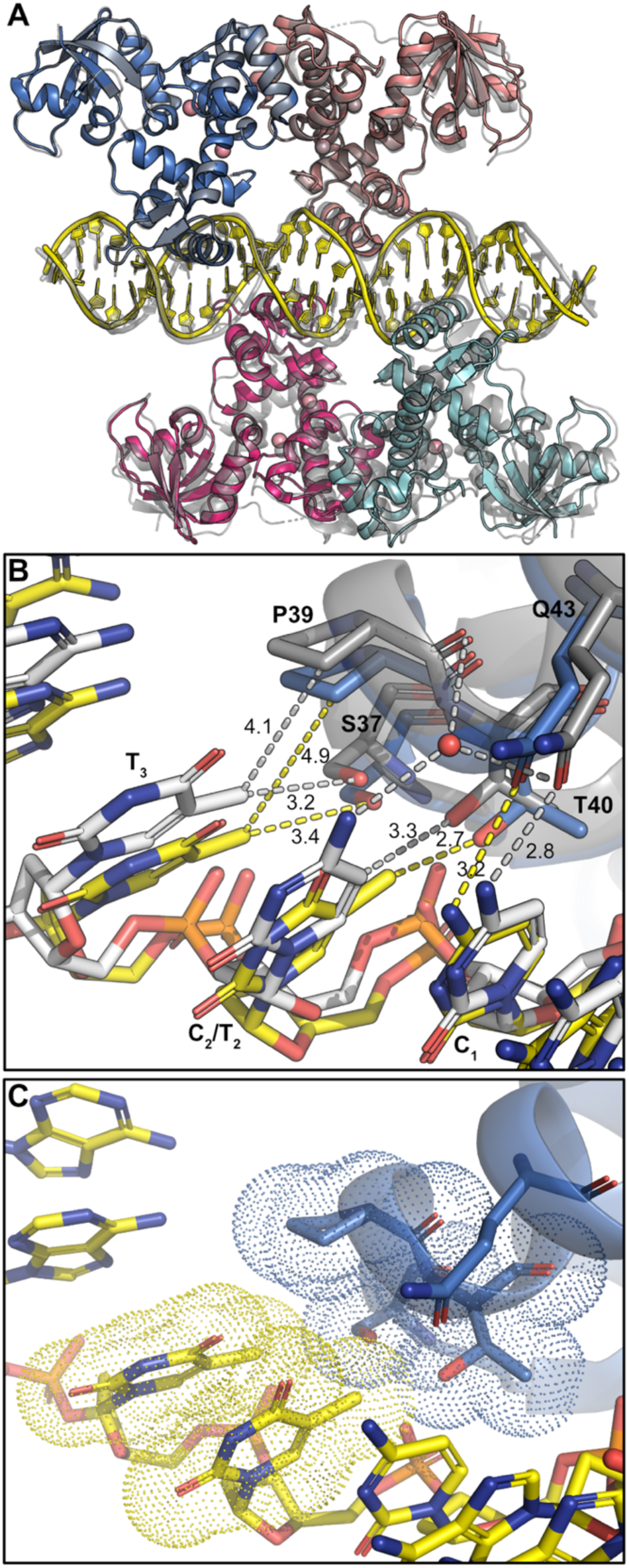
Crystal structure of Co^2+^-activated IdeR^WT^ in complex with a consensus DNA sequence containing the C_2_→T_2_ mutation in every repeat. (**A**) Overall structure of the complex in cartoon representation, superimposed with the structure of the Fe^2+^-activated IdeR^WT^ complex with the full consensus DNA recognition sequence (PDB 7B20) (10). Metal ions are shown as spheres. The complex with the full consensus sequence is shown in transparent grey, while the complex with the T_2_-DNA sequence is colored by subunit (IdeR subunits in marine, salmon, cyan and pink; DNA duplex in yellow). For clarity, only one of the two superimposed DNA chains of the T_2_-DNA complex is shown for each crystallographic symmetry mate making up the physiological complex (see main text). (**B**) Detailed view of the interactions of IdeR subunit A with DNA bases, superimposed with the structure of the IdeR^WT^ complex with the full consensus DNA recognition sequence (PDB 7B20) (10). Sticks are colored by element, with protein and DNA in the T_2_-DNA complex shown with marine and yellow carbon atoms, respectively, and protein and DNA in the full consensus (C_2_) DNA complex with grey and white carbon atoms, respectively. For clarity, only one of the two superimposed DNA chains of the T_2_-DNA complex is shown, and the complementary DNA strands are not shown. Hydrogen bonds and van der Waals interactions are indicated by yellow and grey dashed lines for the T_2_-DNA and full consensus complexes, respectively. Selected interaction distances are labelled in Å. (**C**) Detailed view of the van der Waals interactions of IdeR subunit A with DNA bases in the T_2_-DNA complex, indicated by the dotted van der Waals surfaces of IdeR residues Ser37, Pro39 and Thr40 as well as the DNA bases T_2_ and T_3_. Color scheme as in panel B. Alt text: Crystal structure of Co^2+^-activated IdeR bound to C_1_T_2_T_3_-mutated DNA, compared with the complex with the full consensus DNA, showing overall structural alignment, DNA interactions, hydrogen bonds, and van der Waals contacts.

It is important to note that the C_2_→T_2_ mutation does not disrupt the direct protein–DNA interactions (Fig. 5B and C). The water-mediated interaction between Gln43, Pro39 and the position 2 base is likely to be maintained in the T_2_-DNA complex. While a water molecule was not clearly visible in the electron density and was therefore ultimately not modelled, weak positive difference density in subunit A suggests that it is present at least at partial occupancy. As previously noted in the structure of an IdeR complex with a natural promoter sequence containing a thymine in position 2 of one repeat (10), the methyl group of the thymine in position 2 requires an increased distance between Thr40 and the position 2 nucleotide, compared to the interaction of Thr40 with a cytosine in this position (C_2_-DNA complex: Thr40-Cα – C_2_-C1’ distance 8.2 Å; T_2_-DNA complex: Thr40-Cα – T_2_-C1’ distance 9.1 Å; Fig. 5B and C). This adjustment in turn appears to cause small adjustments in the relative positions of the surrounding recognition helix residues and nucleotides, leading to the other interaction distances between IdeR and DNA bases to be slightly longer in the T_2_-DNA complex than in the full consensus complex. Nevertheless, IdeR forms the same interactions with this mutated sequence as with the full consensus, demonstrating a degree of plasticity in the interaction that is likely necessary for IdeR to function *in vivo*, where no single recognition sequence is perfectly conserved. The structural similarity between non-cognate T_2_-DNA and the cognate C_2_-DNA in complex with IdeR accounts for the weak discrimination of ΔΔG ∼0.5 kcal/mol in our calculations.

If mutations of either the first or the second position of the target site repeat are tolerated by IdeR, then why is a simultaneous mutation of both positions not? The polar side-chain residues Ser37, Gln36, Thr40 and Gln43 of the IdeR recognition helix create a polar environment at the C-H edge of the C-G base pairs in the first and second position of the consensus DNA repeat (Fig. 1B). Additionally, stable non-specific interactions were observed between the phosphate backbone of the consensus sequence (C_1_C_2_T_3_) and the wild-type IdeR protein in the MD trajectories (Supplementary Fig. S1) and crystal structures. Specifically, the phosphate backbone of C_1_ interacts with the side-chains of Gln36 and Thr7, while the C_2_ phosphate engages with the main-chain and side-chain of Ser37 and Thr40, respectively (Fig. 1B). Two non-polar methyl groups of T_1_T_2_ at the C-H edge are strongly disfavored by the polar residues of the recognition helix of IdeR. This increases the free energy of double mutant T_1_T_2_-DNA in the IdeR complex, disrupting the interactions between the phosphate of T_1_ and the protein (Gln36, Thr7) at the first position, as shown in Fig. 6. Consequently, the DNA moves away from the protein, leading to a noticeable narrowing of the major groove. Clearly, a double mutation (C_1_C_2_→T_1_T_2_) causes substantial alterations in the shape and width of the major groove of the DNA within the IdeR complex. The N-terminus of IdeR changed conformation due to the C_1_C_2_→T_1_T_2_ mutation, becoming helical in the T_1_T_2_T_3_-DNA complex, unlike its unstructured state with consensus C_1_C_2_T_3_-DNA, which is required to allow the protein to fit into the target DNA’s major groove (45).

**Figure 6.**
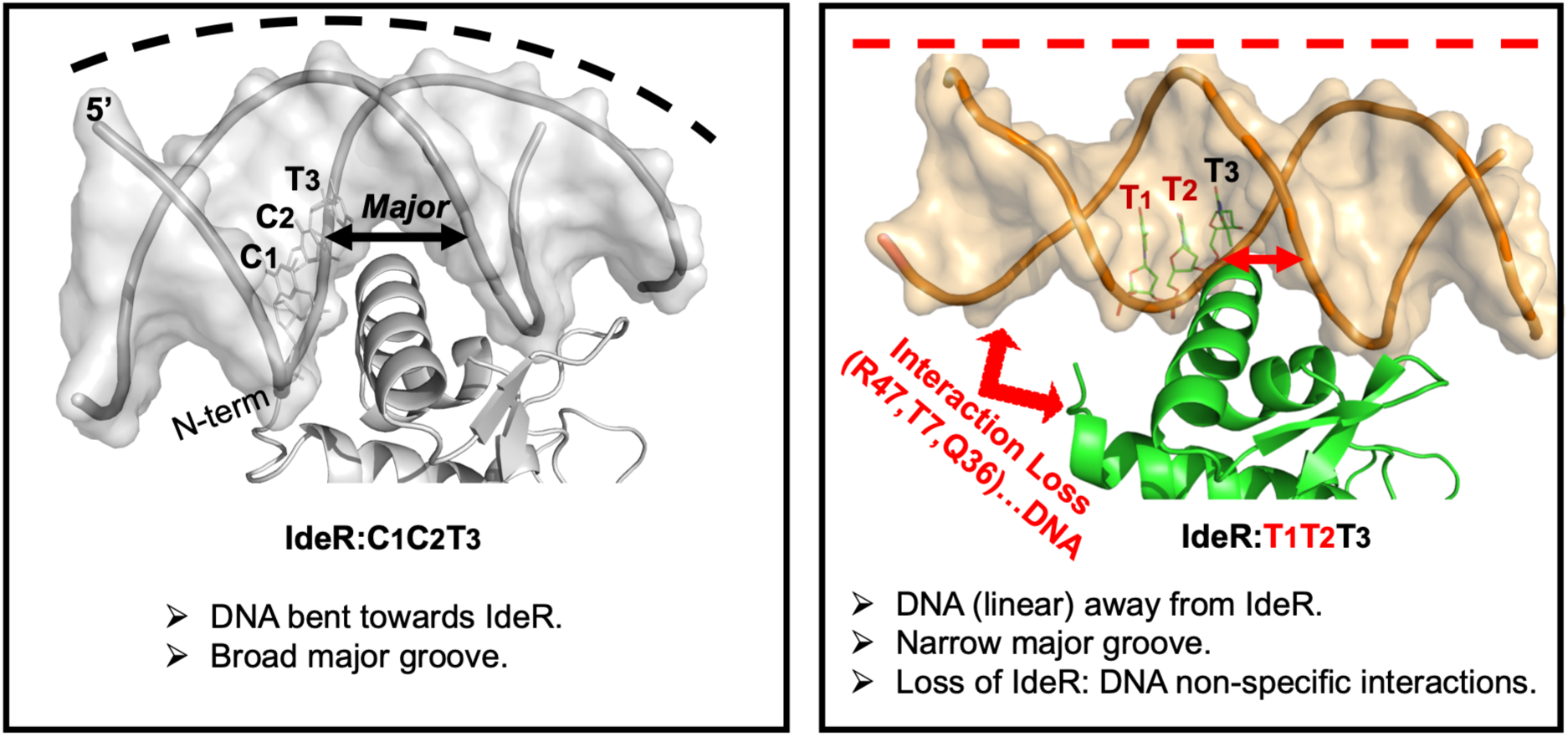
The double C (left) to T (right) mutation in the IdeR–DNA complex significantly alters the shape of the DNA and the size of the groove that accommodates the recognition helix of IdeR. As a result of this double mutation, the non-specific interactions between IdeR and DNA are lost (right). Alt text: Structural comparison showing that the sequence containing the C_1_C_2_→T_1_T_2_ double mutation alters DNA bending and major-groove width, positioning DNA away from IdeR and disrupting non-specific protein–DNA interactions.

Notably, as shown above, a single thymine mutation is allowed at either the first or second position and does not alter the structure of the complex significantly, whereas simultaneous mutation of both positions has a substantial impact on the structure, to the extent that a stable complex cannot be formed. Thus, a hydrophilic-hydrophobic balance in the C-H edge of the first and second base is crucial for stabilizing the IdeR–DNA complex. Increased hydrophobicity beyond the threshold resulted in weakened protein–DNA electrostatic interactions and can penalize the affinity by as much as 4 kcal/mol for T_1_T_2_-DNA.

## DISCUSSION

Previous experimental studies have demonstrated that IdeR/DtxR family iron-dependent regulators from different species can recognize their target 5’-C_1_C_2_T_3_ sequence on the DNA without forming nucleotide-specific interactions (10, 16, 21). It was established that the recognition helix of IdeR positioned in the major groove of the C_1_C_2_T_3_ consensus sequence ensured precise nucleotide recognition. However, the energetics involved in this recognition and its relationship to the protein–nucleotide structure were not understood. Thus, the role of the recognition helix of IdeR in ensuring DNA-binding fidelity remains elusive. We quantitatively assessed the energetics of nucleotide recognition at all three positions of the CCT sequence by the IdeR protein, revealing the principle behind IdeR’s specificity. Using MD simulations and experiments, we demonstrated that a significant free-energy penalty prevents IdeR from misreading non-cognate sequences.

We found that the origin of selectivity for the third nucleotide position (T_3_) is fundamentally different from that of the first two positions (C_1_C_2_). The recognition helix creates a non-polar environment at the C-H edge of the third base (T_3_), particularly involving the Pro39 residue, which provides a hydrophobic cushion for the methyl group of the third thymine base. As a result, the increased polarity from the T_3_→C_3_ mutation near the hydrophobic Pro39 creates strain in the DNA and destabilizes the protein–DNA complex by 3 kcal/mol (Fig. 2). The altered DNA backbone in response to the T_3_→C_3_ mutation was evident in the MD structure (Fig. 4).

In contrast, the recognition helix creates a polar patch at the C-H edge of the first two bases (C_1_ and C_2_), primarily involving four residues: Ser37, Gln36, Thr40 and Gln43. Therefore, a double non-polar substitution (C_1_C_2_ → T_1_T_2_) in the polar patch of the recognition helix of IdeR is highly detrimental, leading to the loss of critical protein (Gln36, Thr7) and phosphate (T_1_) interactions, as well as a relaxation of the DNA backbone into a relatively less compact conformation (Fig. 6). The final structure of the non-cognate T_1_T_2_– IdeR complex was estimated to be destabilized by 4 kcal/mol relative to the cognate C_1_C_2_–IdeR complex. The loosely bound complex with high free energy can drive dissociation of the non-cognate complex, resulting in non-productive binding and preventing misreading.

It was interesting to note that a single cytosine→thymine mutation in either of the first two positions is tolerated by IdeR (i.e, the structure and energetics remained largely unchanged upon a single mutation), whereas mutation of both cytosines is not, demonstrating the importance of the hydrophilic-hydrophobic balance in stabilizing protein–nucleotide interactions. The differential local dielectric created by the recognition helix of IdeR in the C-H edge of the C_1_C_2_ (polar high dielectric) and T_3_ (non-polar low dielectric) bases ensures accurate DNA decoding by imposing strain on the non-cognate sequences, thus destabilizing non-cognate complexes, while allowing some flexibility in recognition of only partially conserved target sequences.

Our finding that Gln43 is dispensable for target DNA recognition by IdeR was initially puzzling, given not only that it is the only residue forming a hydrogen bond with the target DNA, but also its high degree of conservation in the IdeR/DtxR family. In light of the results presented here, it appears likely that its function in DNA binding is the part it plays in creating the polar pocket for the first two positions of the DNA repeat. Just as single mutations in the first two positions of the repeat are tolerated, the mutation of a single IdeR residue that interacts with these positions can be tolerated, whereas larger alterations in the polarity of this pocket most likely are not.

While we did not assess the role of the less conserved positions 4 and 5 of the C_1_C_2_T_3_A_4_A_5_ repeat in IdeR–DNA recognition in this work, our previous work suggests that the adenine bases play a role in recognition although IdeR makes no direct contacts with the last two positions of the repeat (10). We showed that NCTNN and CNTNN repeats are not recognized by IdeR, whereas NCTAA and CNTAA repeats are, and a CCTSS repeat is recognized with lower affinity, indicating that the identity of the bases in these positions is important in the context of the degree of conservation of positions 1-3. Together with the data presented here, this suggests that the adenine bases support recognition by favoring the DNA structure that is compatible with IdeR binding.

Calculated energetics (ΔΔG) provide insights into the fidelity of DNA recognition by IdeR and explain electrophoretic mobility shift assays assessing its binding affinity to various DNA sequences. The simulations demonstrate the versatility of C_1_C_2_T_3_ reading by the IdeR protein, extending well beyond simple protein–DNA interactions. The IdeR protein recognizes its target sequence without engaging in nucleotide-specific interactions by applying stress to the non-target DNA backbone. The recognition helix of IdeR forms a favorable cleft to accommodate the C-H edge of the first three positions of the target sequence (5’-CCT), featuring a polar pocket for the first two bases and a non-polar pocket for the third base. The local dielectric environment destabilizes non-target DNA complexes, distorting the complex and ultimately resulting in non-productive binding. The recognition helix residues of IdeR/DtxR family iron-dependent regulators are highly conserved and recognize the same consensus sequence (10, 14–16), suggesting that they share this DNA recognition mechanism.

## DATA AVAILABILITY

Atomic coordinates and structure factors for the reported crystal structure have been deposited with the Protein Data Bank under accession number 9SUO. The associated raw X-ray diffraction data and all processing files and logs have been deposited at Zenodo under doi: https://doi.org/10.5281/zenodo.23083122. All original EMSA images generated in this study are available at Figshare under doi: 10.6084/m9.figshare.30265090. Input files, topologies, final MD structures, and Q43movie.mpg are provided at Zenodo under doi: https://doi.org/10.5281/zenodo.22951797. GROMACS 2019 (https://manual.gromacs.org/2019/index.html) was used for MD simulations and free energy calculations. The hybrid structures and topologies were generated using the open-source PMX package (https://github.com/deGrootLab/pmx). The open-source version (free for academic users) of PyMOL (https://github.com/schrodinger/pymol-open-source) and VMD 1.9.3 (https://www.ks.uiuc.edu/Research/vmd/) are used for structure generation, visualisation, and trajectory analysis. Analysis of trajectories, structures, and energetics is available in the Supporting Information.

## SUPPLEMENTARY DATA

Supplementary Data are available.

## ACKNOWLEDGEMENTS

PARAM-ISHAN and PARAM-KAMRUPA at IIT Guwahati are gratefully acknowledged for providing the computing facility. The authors would like to thank Diamond Light Source for beamtime (proposal mx37715), and the staff of beamline I24 for assistance with X-ray data collection.

## Author contributions

S.G. performed the computational work. D.M. and S.T. performed protein production and purification. O.K. and J.J.G. performed DNA-binding assays. J.J.G. performed crystallization and structure determination. J.J.G. and P.S. designed the study. S.G., J.J.G. and P.S. analyzed the data and wrote the manuscript with contributions from all authors.

