## Supplementary Information for "The bacterial iron sensor IdeR distinguishes DNA sequences by applying stress to non-cognate DNA"

### Contents

#### Supplementary Figures

**Figure S1.** Structural and dynamical parameters from the MD trajectory.

**Figure S2:** Additional EMSA analysis of $IdeR^{WT}$ binding to the Consensus_$T_3$-$C_3$ DNA sequence.

**Figure S3:** Superposition of the crystal structures of $Co^{2+}$-activated $IdeR^{WT}$ in complex with a consensus DNA sequence containing the $C_2 \rightarrow T_2$ mutation in every repeat and the $Fe^{2+}$-activated $IdeR^{WT}$ complex with the full consensus DNA recognition sequence.

#### Supplementary Tables

**Table S1**: Calculated free energies.

**Table S2:** List of oligonucleotides used in this study.

#### Supplementary References

### Supplementary Figures

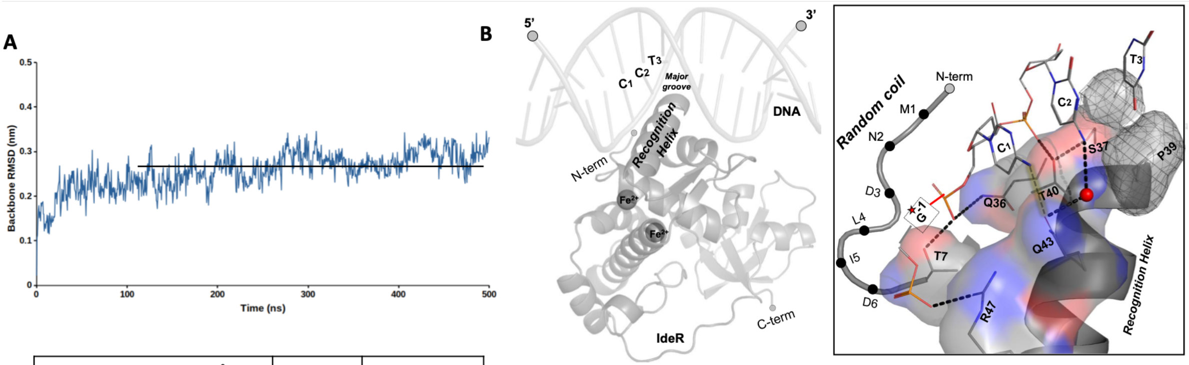

C

| Distances (in Å) | X-ray | MD |
|---|---|---|
| R47(NH1)-$G^{-1}$(O2P) | 2.6 | 4.5 ± 0.4 |
| T7(OG1)-$C_1$(O1P) | 2.6 | 3.1 ± 0.4 |
| Q36(OE1)-$C_1$(O2P) | 3.2 | 3.4 ± 0.2 |
| T40(OG1)-$C_2$(O2P) | 2.5 | 2.8 ± 0.1 |
| S37(OG)-$C_2$(O1P) | 3.1 | 2.7 ± 0.1 |
| Q43(OG1)-$C_1$(N4) | 3.1 | 8.5 ± 0.3 |
| Q43(OG1)-$C_2$(N4) | 4.7 | 7.5 ± 0.3 |

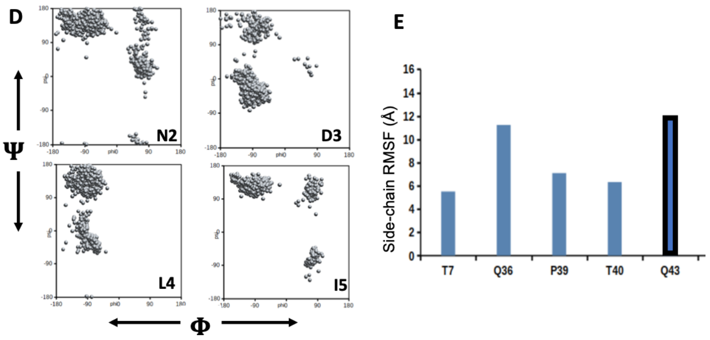

**Figure S1.** Structural and dynamical parameters from the MD trajectory. (**A**) Backbone RMSD of IdeR as a function of time from the cognate complex (PDB 7B20) (1). (**B**) The recognition helix of wild-type IdeR is placed in the major groove of the cognate DNA. The zoomed-in view shows the protein-DNA interaction network (dashed lines). The Pro39 residue provides a hydrophobic cushion for the methyl group of $T_3$ in the cognate $C_1C_2T_3$ DNA, as shown in the surface mesh. (**C**) Comparison of interatomic distances between the crystal structure and MD trajectory. The standard deviation is reported as an error. (**D**) Ramachandran plot of four residues from the N-terminal of IdeR. (**E**) Side-chain RMSF of selected residues in the protein-DNA interface.

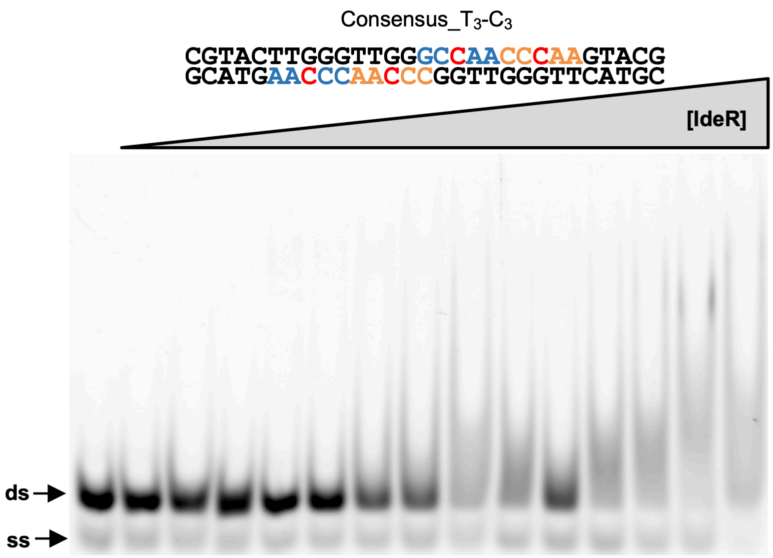

**Figure S2.** Additional EMSA analysis of IdeR$^{WT}$ binding to the Consensus_$T_3$-$C_3$ DNA sequence. IdeR was added in increasing concentrations (150 nM – 22.5 µM dimer) to 30 nM fluorescence-labeled double-stranded DNA probe in the presence of 100 µM $Co^{2+}$ and competitor DNA. Note that the IdeR concentrations used here are 10-fold higher compared to Figure 3. Protein-DNA complexes were resolved on a 4% Tris-acetate polyacrylamide gel. The left-most lane is a control reaction without protein. ds, unbound double-stranded DNA probe; ss, non-hybridized single-stranded DNA.

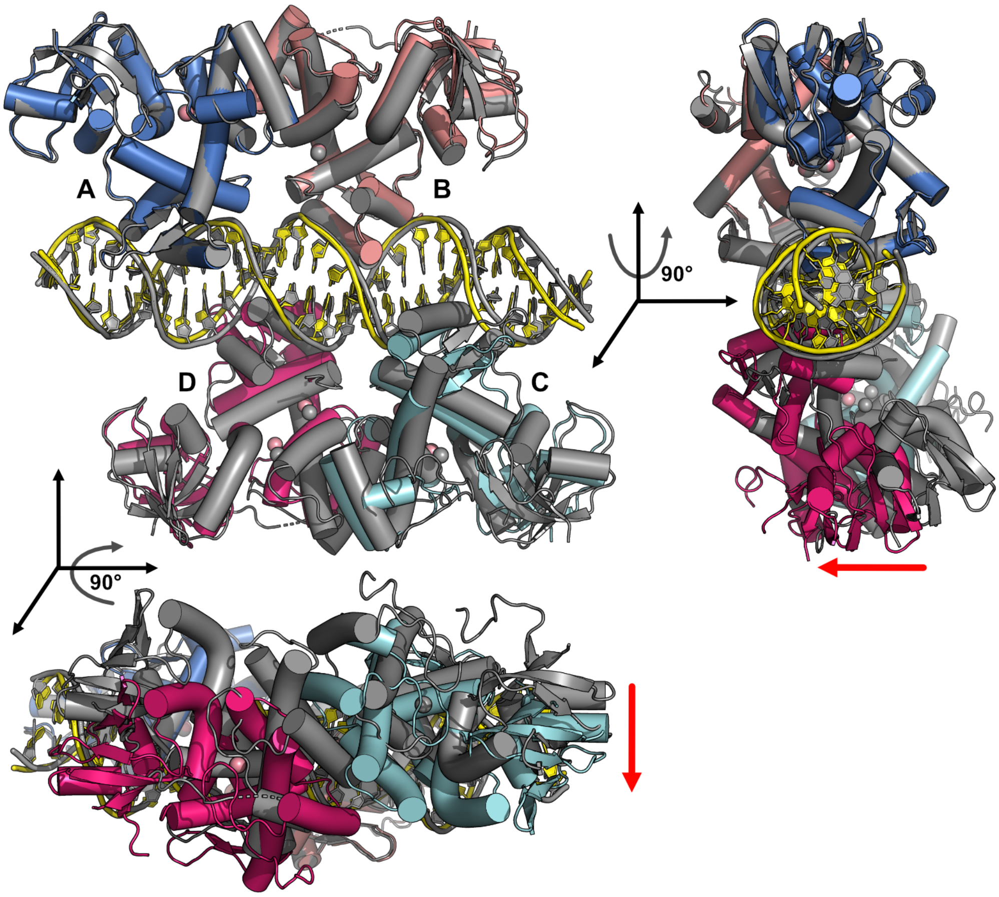

**Figure S3.** Superposition of the crystal structures of $Co^{2+}$-activated $IdeR^{WT}$ in complex with a consensus DNA sequence containing the $C_2 \rightarrow T_2$ mutation in every repeat (colored by subunit), and the $Fe^{2+}$-activated $IdeR^{WT}$ complex with the full consensus DNA recognition sequence (PDB 7B20; shown in grey) (1). Three orthogonal views illustrate the shift of one IdeR dimer (chains C and D) in the $T_2$-DNA complex in comparison to the full consensus ($C_2$) DNA complex. Red arrows indicate the direction of the shift.

### Supplementary Tables

**Table S1**. Calculated free energies.

| Alchemical free energy ($\Delta G^{comp}$, $\Delta G^{free}$ in kcal/mol) | | | | |
|---|---|---|---|---|
| Alchemical Transformation | Replicas | $\Delta G^{comp}$ | $\Delta G^{free}$ | Average $\Delta\Delta G = \Delta G^{comp} - \Delta G^{free}$ |
| $C_1 \rightarrow T_1$ | 1 | 134.04 ± 0.27 | 134.19 ± 0.15 | -0.15 ± 0.05 |
| | 2 | 134.01 ± 0.24 | 134.12 ± 0.12 | |
| | 3 | 134.07 ± 0.27 | 134.27 ± 0.14 | |
| $C_2 \rightarrow T_2$ | 1 | 133.78 ± 0.12 | 134.08 ± 0.04 | -0.34 ± 0.05 |
| | 2 | 133.80 ± 0.14 | 134.13 ± 0.07 | |
| | 3 | 133.72 ± 0.15 | 134.12 ± 0.05 | |
| $C_1C_2 \rightarrow T_1T_2$ | 1 | 272.28 ± 0.35 | 268.26 ± 0.23 | +4.11 ± 0.10 |
| | 2 | 272.38 ± 0.31 | 268.29 ± 0.27 | |
| | 3 | 272.42 ± 0.27 | 268.21 ± 0.20 | |
| $T_3 \rightarrow C_3$ | 1 | -131.44 ± 0.20 | -134.05 ± 0.09 | +2.68 ± 0.09 |
| | 2 | -131.34 ± 0.20 | -134.12 ± 0.05 | |
| | 3 | -131.46 ± 0.19 | -134.10 ± 0.10 | |
| Q43→ A43 | 1 | 55.02 ± 0.13 | 55.15 ± 0.25 | -0.13 ± 0.01 |
| | 2 | 55.15 ± 0.17 | 55.27 ± 0.20 | |
| | 3 | 55.12 ± 0.14 | 55.26 ± 0.23 | |
| MM/PBSA binding affinity ($\Delta G$ in kcal/mol) | | | | |
| | | $\varepsilon = 1$ | $\varepsilon = 5$ | $\Delta\Delta G = \Delta G(G39) - \Delta G(P39)$ |
| Wild-type (P39) | $\Delta G$ | -5.06 ± 0.97 | -4.67 ± 0.84 | 4.10 ± 0.78 ($\varepsilon = 1$)<br>2.82 ± 0.82 ($\varepsilon = 5$) |
| Mutant (G39) | $\Delta G$ | -0.96 ± 0.31 | -1.68 ± 0.35 | |

**Table S2**. List of oligonucleotides used in this study.

| Name | Sequence 5'→3' | Description |
|---|---|---|
| Cons_FAM_fwd | [FAM]-CGTACTTAGGTTAGGCTAACCTAAGTACG | Forward strand of the Consensus_Full sequence used for EMSA analyses (5' FAM label) |
| Cons_rev | CGTACTTAGGTTAGCCTAACCTAAGTACG | Reverse strand of the Consensus_Full sequence used for EMSA analyses |
| Cons_T3-C3_FAM_fwd | [FAM]-CGTACTTGGGTTGGGCCAACCCAAGTACG | Forward strand of the Consensus_$T_3$-$C_3$ sequence used for EMSA analyses (5' FAM label) |
| Cons_T3-C3_rev | CGTACTTGGGTTGGCCCAACCCAAGTACG | Reverse strand of the Consensus_$T_3$-$C_3$ sequence used for EMSA analyses |
| Cons_C2-T2_fwd | CGTACTTAAGTTAAGTTAACTTAAGTCACG | Forward strand of the Consensus_$C_2$-$T_2$ sequence used for crystallization |
| Cons_C2-T2_rev | CGTGACTTAAGTTAACTTAACTTAAGTACG | Reverse strand of the Consensus_$C_2$-$T_2$ sequence used for crystallization |
